# Genes Encoding ACE2, TMPRSS2 and Related Proteins Mediating SARS-CoV-2 Viral Entry are Upregulated with Age in Human Cardiomyocytes

**DOI:** 10.1101/2020.07.07.191429

**Authors:** Emma L. Robinson, Kanar Alkass, Olaf Bergmann, Janet J. Maguire, H. Llewelyn Roderick, Anthony P. Davenport

**Affiliations:** Department of Cardiology, Cardiovascular Research Institute Maastricht (CARIM) Maastricht University, The Netherlands and Laboratory of Experimental Cardiology, Dept. of Cardiovascular Sciences, KU Leuven, Campus Gasthuisberg, Herestraat 49, B-3000, Leuven, Belgium; Karolinska Institutet, BioClinicum, Oncology and Pathology, SE-17177 Stockholm, Sweden; The National Board of Forensic Medicine, Stockholm, Sweden; Center for Regenerative Therapies Dresden, TU-Dresden, Fetscherstrasse 105, 01307 Dresden, Germany; Karolinska Institutet, Biomedicum, Cell and Molecular Biology, SE-17177 Stockholm, Sweden; Experimental Medicine and Immunotherapeutics, University of Cambridge, Addenbrooke’s Hospital, Cambridge U.K; KU Leuven, Department of Cardiovascular Sciences, Laboratory of Experimental Cardiology, 3000 Leuven, Belgium and K.G. Jebsen Center for Cardiac Research, University of Oslo, Oslo, Norway

**Keywords:** Human Cardiomyocytes, RNA-sequencing, Angiotensin, Apelin, Bradykinin, SARS-CoV-2, COVID-19

## Abstract

Age is an independent risk factor for adverse outcome in patients following COVID-19 infection. We hypothesised that differential expression of genes encoding proteins proposed to be required for entry of SARS-Cov-2 in aged compared to younger cardiomyocytes might contribute to the susceptibility of older individuals to COVID-19-associated cardiovascular complications.

We generated strand-specific RNA-sequencing libraries from RNA isolated from flow-sorted cardiomyocyte nuclei from left ventricular tissue. RNASeq data were compared between five young (19-25yr) and five older (63-78yr) Caucasian males who had not been on medication or exhibited evidence of cardiovascular disease post-mortem.

Expression of relevant genes encoding *ACE2, TMPRSS2, TMPRS11D, TMPRS11E, FURIN, CTSL, CTSB* and *B^0^AT1/SLC6A19* were upregulated in aged cardiomyocytes and the combined relative cardiomyocyte expression of these genes correlated positively with age. Genes encoding proteins in the RAAS and interferon/interleukin pathways were also upregulated such as *ACE, AGTR1, MAS1* and *IL6R*.

Our results highlight SARS-CoV-2 related genes that have higher expression in aged compared with young adult cardiomyocytes. These data may inform studies using selective enzyme inhibitors/antagonists, available as experimental compounds or clinically approved drugs e.g. remdesivir that has recently been rapidly accepted for compassionate use, to further understand the contribution of these pathways in human cardiomyocytes to disease outcome in COVID-19 patients.

---

Age (>70 years), case fatality rate (CFR,10.2%) and coexisting conditions, particularly cardiovascular disease (CFR,10.5%) and hypertension (CFR,6.0%), are independent predictors of adverse outcome for 45,000 COVID-19 patients in China^1^. A consensus has emerged that SARS-CoV-2 uses the same ‘receptor’ as SARS-CoV, the angiotensin converting enzyme 2 (ACE2), for initial binding to the host cell. This must be co-expressed with the serine protease TMPRSS2, that primes the spike protein S1 for endocytosis-mediated internalization of virus, employing the S2 domain for fusion to the host membrane (Figure [1A])^2-4^. A key difference in SARS-CoV-2 is a spike protein second site (S2’), proposed to be cleaved by the proteinase furin^2^. Once inside the cell cysteine proteases, cathepsin L and B, are thought critical for endosomal processing in certain cells^3,4^.

**Figure 1.**
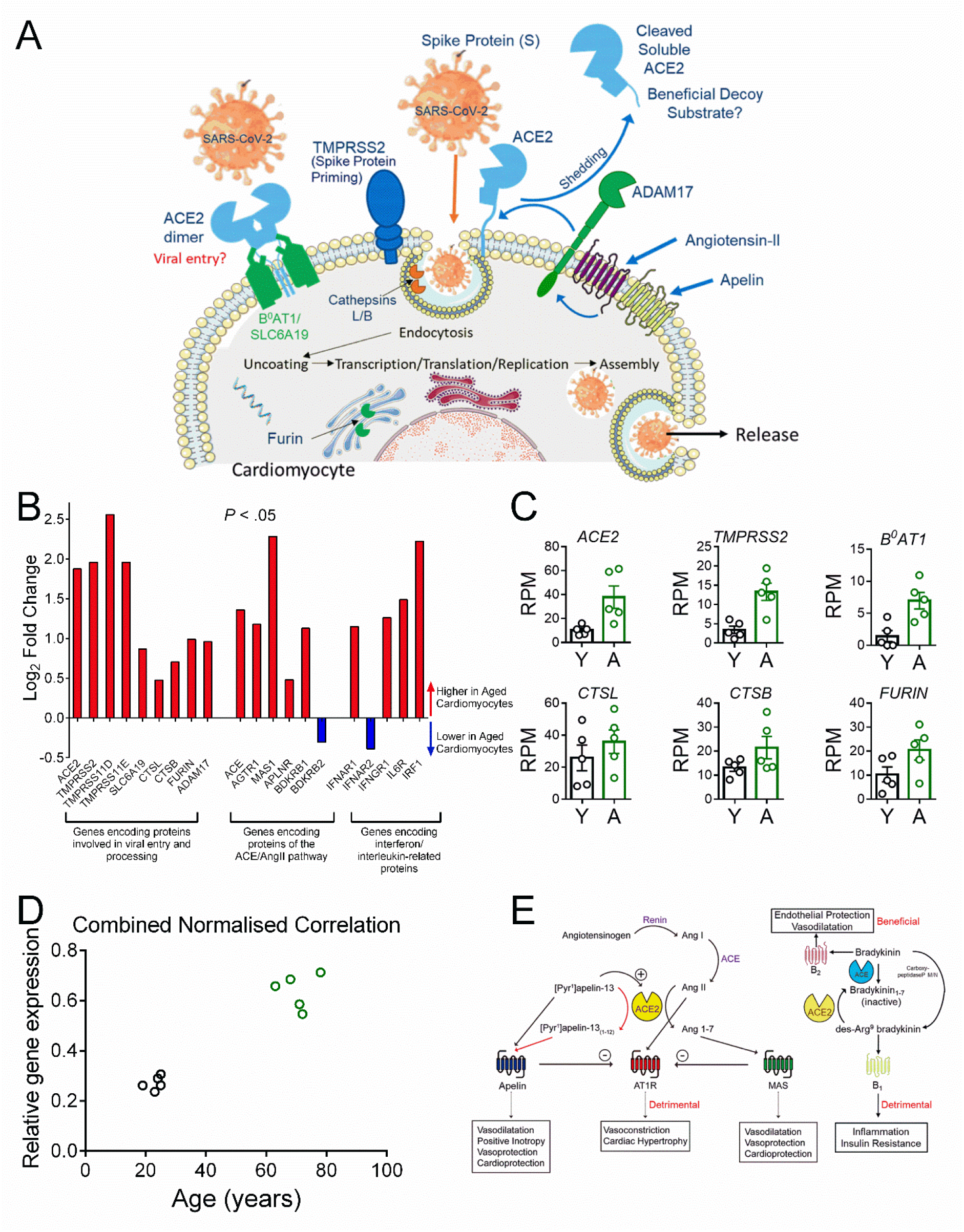
Genes Encoding Proteins Mediating Viral Entry are Upregulated with Age. References are limited to ten but see recent review^5^ for a comprehensive list of references supporting the concepts outlined in this letter. **[A]** Schematic diagram of the key proteins predicted from RNASeq data to be expressed by human cardiomyocytes. We propose SARS-CoV-2 binds initially to ACE2 (with the ACE2/B^0^AT1 complex as a potential second entry site). TMPRSS2 priming of the spike protein S1 together with further protease activation by Cathepsins B and L facilitate viral cell entry and internalization by endocytosis. Furin may also have a role in this process. Internalization of the virus with ACE2 inhibits ACE2 carboxypeptidase activity that normally hydrolyses Ang-II, [Pyr^1^]-apelin-13 and des-Arg^9^-bradykinin. ADAM17, present on the cell surface, cleaves ACE2 to the soluble form that circulates in the plasma and could act as a decoy substrate for the virus. Levels of ADAM17 may be regulated by Ang-II and apelin acting via their respective G-protein coupled receptors^4^. **[B]** Log_2_ Fold change for expressed genes encoding SARS-CoV-2 entry proteins, peptide receptors in ACE/ACE2 pathway and receptors for IL-6 and interferon in aged cardiomyocytes. Strand-specific RNA-sequencing libraries were generated from RNA isolated from flow-sorted cardiomyocyte nuclei from left ventricular tissue^10^. RNASeq data were compared between five young (19-25yr) and five older (63-78yr) Caucasian males, not on medication, with no evidence of cardiovascular disease post-mortem. A Chi Squared analysis for the selected gene panel shown in [B] showed significant enrichment for differentially expressed genes compared with the RNASeq data set as an entirety (*P* < .05, 95 % confidence level). *ACE2*, angiotensin I converting enzyme 2; *TMPRSS2*, transmembrane protease, serine *2;TMPRSS11D*, transmembrane protease, serine 11D; *TMPRSS11E*, transmembrane protease, serine 11E; *FURIN*, furin (paired basic amino acid cleaving enzyme); *CTSB*, cathepsin B; *CTSC*, cathepsin C; *CTSF*, cathepsin F; *CTSG*, cathepsin G; *CTSK*, cathepsin K; *CTSL;* cathepsin L; *CTSO*, cathepsin O; *CTSS*, cathepsin S; *CTSV*, cathepsin V; *CTSZ*, cathepsin Z; *SLC6A19*, solute carrier family 6 (neutral amino acid transporter), member 19; *ADAM17*, ADAM metallopeptidase domain 17; *ACE*, angiotensin I converting enzyme; *AGTR1;* angiotensin II receptor, type 1; *MAS1*, MAS1 proto-oncogene, G protein-coupled receptor; *BDKRB1*, bradykinin receptor B1; *BDKRB2*, bradykinin receptor B2; *APLNR*, apelin receptor; *IFNAR1*, interferon (alpha, beta and omega) receptor 1; *IFNAR2*, interferon (alpha, beta and omega) receptor 2, (*IFNAR1 and 2* form chains for interferons alpha and beta); IFNGR1, *Interferon gamma receptor 1, IL6R*, interleukin 6 receptor; *IRF1*, interferon regulatory factor 1 (regulate transcription of interferons). **[C]** Scatter dot plot showing individual data points (error bars indicate mean±S.E.M) of key genes with higher expression in young (Y) verses aged (A) cardiomyocytes. Cardiomyocytes were double positive for *ACE2* and *TMPRSS2*, critical for viral entry; ACE2 chaperone, B^0^AT1 *(SLC6A19)*, a potential second mechanism for entry; *Cathepsins B, L and FURIN* proposed to mediate further protease activation to facilitate virus entry and internalization **[D]** Cardiomyocyte expression levels of the gene panel shown in [C] correlates positively with age. For each of the six targets presented in [C], the mean relative gene expression was calculated, normalized to the sample with the highest expression in CPM (non-parametric Spearman correlation, two-tailed, r = 0.86, *P* =0.0025). **[E]** Schematic diagram of the genes encoding GPCRs that are expressed and higher in expression in aged human cardiomyocytes compared with young. Activation of the renin-angiotensin system, with overproductuon of Ang-II, contributes to acute respiratory distress syndrome following infection by coronoviruses. The Ang-II synthetic enzyme ACE and the ANG-II cognate receptor gene *AGTR1* increased with age, together with B1 receptor gene, *BDKBR1*. This bradykinin receptoris selectively activated by des-Arg^9^-bradykinin (that would normally be inactiveated by ACE2), representing a second deleterious pathway. Both pathways could be blocked with clinically approved drugs (ACE inhibitors, AT1 receptor antagonists and the B2 receptor antagonist Icatibant) for secondary treatment in pateints with COVID-19. Internalization of ACE2 by the virus is predicted to increase Ang II levels but reduce those of Ang1-7 but the gene encoding the proposed Ang1-7 receptor, *MAS*, was still detected in aged myocytes, as was the apelin receptor. The beneficial potent positive inotropic action of apelin in the heart as well as its anti-thrombotic and anti-diabetic properties, suggests its receptor is a promising therapeutic target for administered apelin and this strategy is currently under clinical investigation.

In the cardiovascular system, ACE2 protein^5^ and mRNA^6^ are present in cardiomyocytes. ACE2 normally functions as a carboxypeptidase cleaving single C-terminal amino acids thus hydrolysing Pro-Phe in Ang-II to Ang1-7, [Pyr^1^]-apelin-13 to [Pyr^1^]-apelin(1-12) and des-Arg^9^-bradykinin to inactive bradykinin(1-8)^4^. Internalization of ACE2 by virus potentially reduces the beneficial counter regulatory function of these peptide products to the RAAS pathway^2,4^. Conversely, the serine protease ADAM17 cleaves ACE2 to release its ectodomain and this is stimulated by Ang-II and, potentially, apelin acting via their respective G-protein coupled receptors^7^. Shed ACE2 binds SARS-CoV-2, a complex predicted not to internalize, and therefore circulating ACE2 could be exploited as a beneficial viral decoy substrate. Intriguingly, ACE2 is highly expressed in the GI tract where it is associated with B^0^AT1 (SLC6A19) that actively transports most neutral amino acids across the apical membrane of epithelial cells^4,8^. It is not yet known if B^0^AT1 and ACE2 are coexpressed in cardiomyocytes and represent an important mechanism of viral entry, but they can form a heterodimer, with the ACE2 capable of binding the spike protein S1^8^. Interleukin 6 (IL-6), normally transiently produced, is elevated in serum and positively correlated with disease severity in COVID-19 patients^9^.

We hypothesised that differential expression of genes encoding proteins in these pathways in aged cardiomyocytes could explain why the myocardium of older patients might be particularly vulnerable to the virus, manifesting as cardiovascular complications such as myocarditis. To selectively analyse cardiomyocyte gene expression, we generated strand-specific RNA-sequencing libraries from RNA isolated from flow-sorted cardiomyocyte nuclei from left ventricular tissue^10^. RNASeq data were compared between five young (19-25yr) and five older (63-78yr) Caucasian males, not on medication, with no evidence of cardiovascular disease post-mortem.

## Viral entry, membrane fusion and endocytosis

The key finding (Figure [1B], [1C]) is that expression of genes encoding proteins hypothesised as important for viral entry, crucially including *ACE2* and *TMPRSS2*, were upregulated in aged cardiomyocytes, as well as *TMPRS11E* and *TMPRS11D*, (where function is less well established), *FURIN* and cathepsins *CTSL* and *CTSB*. Importantly, while *B^0^AT1/SLC6A19* was minimally expressed in young cardiomyocytes it was upregulated in aged cardiomyocytes. Moreover, combined relative cardiomyocyte expression of these genes correlated positively with age (Figure [1D]).

## ACE/ACE2 regulation

The expression of Ang-II synthetic enzyme *ACE* and cognate receptor gene *AGTR1* together with *BDKBR1* increased with age, the latter normally not expressed in myocytes until induced by inflammation and selectively activated by des-Arg^9^-bradykinin (Figure [1B], [1E])^9^. Both are potentially deleterious peptides, that we hypothesise would be increased by loss of cell surface ACE2 caused by viral internalization or ADAM17-mediated shedding. Expression of genes encoding receptors for peptides that normally mediate beneficial counter-regulatory effects to Ang-II in the heart, *MAS1*/Ang1-7, *APLNR*/apelin and *BDKRB2*/bradykinin were both up and downregulated.

## Inflammatory mediators and endogenous antiviral strategies

Penetration of SARS-CoV-2 into lung alveoli, resulting in the ‘cytokine storm’ is a major determinant for COVID-19 patient intubation and mortality. IL-6 is a key mediator^4^: its receptor *(IL6R)* is expressed on cardiomyocytes, with a modest increase with age suggesting anti-IL-6R monoclonal antibodies, currently in trials^10^ may prove cardioprotective.

Unlike most other cells, cardiomyocyte numbers remain stable, regenerating slowly. Although particularly vulnerable to viral infection they have evolved intrinsic mechanisms to combat cytotoxic effects and viral replication mainly by generating type 1 interferon responses. SARS-CoV-2 counterattacks by producing proteins that interfere with interferon pathways. While interferon receptor genes *(IFNAR1/IFNAR2)* were expressed in cardiomyocytes, no pattern emerged with age.

Our results highlight SARS-CoV-2 related genes that have higher expression in aged compared with young adult cardiomyocytes. Our objective is to foster confirmation (or rebuttal) by studies on relevant protein levels in cardiomyocytes and using selective enzyme inhibitors/antagonists to dissect these pathways; available as experimental compounds or clinically approved drugs e.g. remdesivir has recently been rapidly accepted for compassionate use (https://www.guidetopharmacology.org/coronavirus.jsp). Drugs to treat both acute and long-term effects of SARS-CoV-2 will need to focus on these three key stages of infection: preventing the virus entering cells, stopping viral replication and reducing resulting tissue damage, particularly in the hearts of aged individuals that are the most vulnerable members of society to COVID-19.

## ACKNOWLEDGEMENTS

Heart samples were obtained from the KI Donatum, Karolinska Institutet, Stockholm, Sweden with ethical approval. Quality control and sequencing was carried out at The Babraham Institute Sequencing Facility, Cambridge (UK).

## References

1. Sommerstein R. Preventing a covid-19 pandemic: ACE inhibitors as a potential risk factor for fatal Covid-19. BMJ. 2020 368: doi: https://doi.org/10.1136/bmj.m810.

2. Liu PP, Blet A, Smyth D, Li H. The Science Underlying COVID-19: Implications for the Cardiovascular System. Circulation. 2020. doi: 10.1161/CIRCULATIONAHA.120.047549.

3. Hoffmann M, Kleine-Weber H, Schroeder S, Kruger N, Herrler T, Erichsen S, Schiergens TS, Herrler G, Wu NH, Nitsche A, Muller MA, Drosten C, Pohlmann S. SARS-CoV-2 Cell Entry Depends on ACE2 and TMPRSS2 and Is Blocked by a Clinically Proven Protease Inhibitor. Cell. 2020;181(2):271–80 e8. Doi:10.1016/j.cell.2020.02.052.

4. Alexander SPH, Armstrong J, Davenport AP, Davies J, Faccenda E, Harding SD, Levi-Schaffer F, Maguire JJ, Pawson AJ, Southan C, Spedding MJ. A rational roadmap for SARS-CoV-2/COVID-19 pharmacotherapeutic research and development. IUPHAR Review 29. Br J Pharmacol. 2020. Doi:10.1111/bph.15094.

5. Hikmet F, Méar L, Edvinsson A, Micke P, Uhlén M, Lindskog C. The protein expression profile of ACE2 in human tissues bioRxiv 2020.03.31.016048; doi: https://doi.org/10.1101/2020.03.31.016048.

6. Tucker NR, Chaffin M, Bedi KC, Papangeli I, Akkad AD, Arduini A, Hayat S, Eraslan G, Muus C, Bhattacharyya R, Stegmann CM, Margulies KB, Ellinor PT. Myocyte Specific Upregulation of ACE2 in Cardiovascular Disease: Implications for SARS-CoV-2 mediated myocarditis. medRxiv. 2020. 10.1101/2020.04.09.20059204.

7. Xu J, Sriramula S, Xia H, Moreno-Walton L, Culicchia F, Domenig O, Poglitsch M, Lazartigues E. Clinical Relevance and Role of Neuronal AT1 Receptors in ADAM17-Mediated ACE2 Shedding in Neurogenic Hypertension. Circ Res. 2017;121(1):43–55. 10.1161/CIRCRESAHA.116.310509.

8. Yan R, Zhang Y, Li Y, Xia L, Guo Y, Zhou Q. Structural basis for the recognition of SARS-CoV-2 by full-length human ACE2. Science. 2020;367(6485):1444–8. 10.1126/science.abb2762.

9. Toniati P, Piva S, Cattalini M, Garrafa E, Regola F, Castelli F, et al Tocilizumab for the treatment of severe COVID-19 pneumonia with hyperinflammatory syndrome and acute respiratory failure: A single center study of 100 patients in Brescia, Italy. Autoimmun Rev. 2020;19, 102568. 10.1016/j.autrev.2020.102568.

10. Thienpont B, Aronsen JM, Robinson EL, Okkenhaug H, Loche E, Ferrini A, Brien P, Alkass K, Tomasso A, Agrawal A, et al. The H3K9 dimethyltransferases EHMT1/2 protect against pathological cardiac hypertrophy. J Clin Invest. 2017 127:335–348. doi: 10.1172/JCI88353.

